# The Maternal Effect of SKN-1B and DAF-7 on Intergenerational Pathogen Avoidance Learning in *C. elegans*

**DOI:** 10.1101/2025.04.14.648668

**Authors:** Rebekka Paisner, Mattias Bekele, Ebubechi Onyia, Andrew Gordus

**Affiliations:** Department of Biology, Johns Hopkins University, Baltimore, MD; Solomon H. Snyder Department of Neuroscience, Johns Hopkins University, Baltimore, MD

## Abstract

Parental health strongly influences the development of embryos, and can ultimately influence behavior post-birth. When exposed to pathogenic *Pseudomonas aeruginosa* (PA14), *C. elegans* produces a robust immune response, as well as a learned behavior to avoid PA14 in the future. This learned avoidance of PA14 can be inherited by the F1 generation. An important trigger of the worm’s immune response to infection is the Nrf2 homologue, SKN-1. We find that the SKN-1B isoform strongly influences the intergenerational inheritance of PA14 avoidance in F1 animals via DAF-7 (TGF-β) signaling from ASI. Functional SKN-1B is required in both the P0 and F1 generations to facilitate F1 avoidance of PA14 due to parental exposure. While inherited avoidance of PA14 was not observed in F2+ generations, the single generation learning investigated here may provide a parallel pathway to improve the fitness of F1 animals in a dynamic environment.

## Introduction

Organisms rely on sensory cues to make adaptive decisions in response to their environment. *Caenorhabditis elegans*, a model nematode, encounters a diverse microbial landscape in which bacteria serve as both food sources and potential threats (1). While the worm exhibits innate attraction to bacterial odors, the safety of consuming these bacteria is highly contextual. Some bacterial species, such as *Pseudomonas aeruginosa* (PA14), present a unique challenge: although nutritious under certain conditions, they can become pathogenic and lethal at higher temperatures (2). This necessitates a learned aversive response, wherein worms associate bacterial cues with prior experiences of infection and stress.

Prior work has demonstrated that exposure to PA14 induces an avoidance behavior in *C. elegans*, as well as an upregulation of the TGF-β homologue *daf-7*, in the ASJ and ASI sensory neurons. Remarkably, this learned aversion is transmitted to offspring in a *daf-7* dependent manner, raising questions about the underlying mechanisms that govern behavioral persistence across generations. However, key aspects of this phenomenon remain unresolved, including the precise neuronal and molecular pathways that enable the transmission of pathogen avoidance to the next generation (3–5).

In addition to a learned avoidance of PA14, pathogen infection triggers a robust immune response to improve fitness. An important trigger for this response is the Nrf2 homologue, SKN-1 (6–9). Several isoforms of this gene exist. While infection induces a strong induction of SKN-1 activity in the gut, neuronal SKN-1 is also important for triggering systemic changes in metabolism (8, 10). Many of these processes rely on the *skn-1* isoform, *skn-1c*. However, the *skn-1b* isoform is specifically expressed in the ASI neurons (11), which also increase expression of *daf-7* upon PA14 infection (3, 5). We find that the increased expression of *daf-7* in ASI neurons relies on a decreased expression of *skn-1b*. While this does not appear to affect learned avoidance of PA14, it does appear to affect the inherited avoidance in the F1 generation. The expression of *skn-1b* in both the parents and the F1 generation is necessary to ensure inherited PA14 avoidance. This effect only lasts for one generation, and may act in parallel to multi-generational inheritance.

## Results

### DAF-7 is required for inherited avoidance of PA14 in F1 animals

Previous findings have shown that after training on PA14 for 24 hours, *C. elegans* have elevated expression of *daf-7* in the ASJ and ASI neurons(3, 12). This increase in *daf-7* expression correlates with a learned avoidance of PA14. Behaviorally, when given the choice between the known beneficial strain of *E. coli*, OP50 and the pathogenic PA14, *C. elegans* that were trained on toxic PA14 will chemotax away from PA14 towards OP50 (1, 3, 13, 14).

Remarkably, this learned avoidance behavior can be passed to subsequent generations, suggesting the involvement of transgenerational memory (3, 5). To confirm the robustness of this paradigm, we entrained *C. elegans* on PA14 for 24 hours (**Fig. 1A**), and observed avoidance of PA14 in both the parental (P0) and first filial generation (F1) (**Fig. 1B**), although we did not see a robust avoidance in subsequent generations (**Fig. S1**). Since the F1 response was a consistent observation, and the conditions for observed avoidance in subsequent generations is debated (4, 5), we limited our experiments to the P0 and F1 generations. Consistent with prior reports (3, 12), we observed elevated expression of the *daf-7p::Venus* reporter in the ASI neurons of PA14-trained P0 animals, as well as in the F1 progeny of PA14-trained worms (**Fig. 1C**)

**Figure 1:**
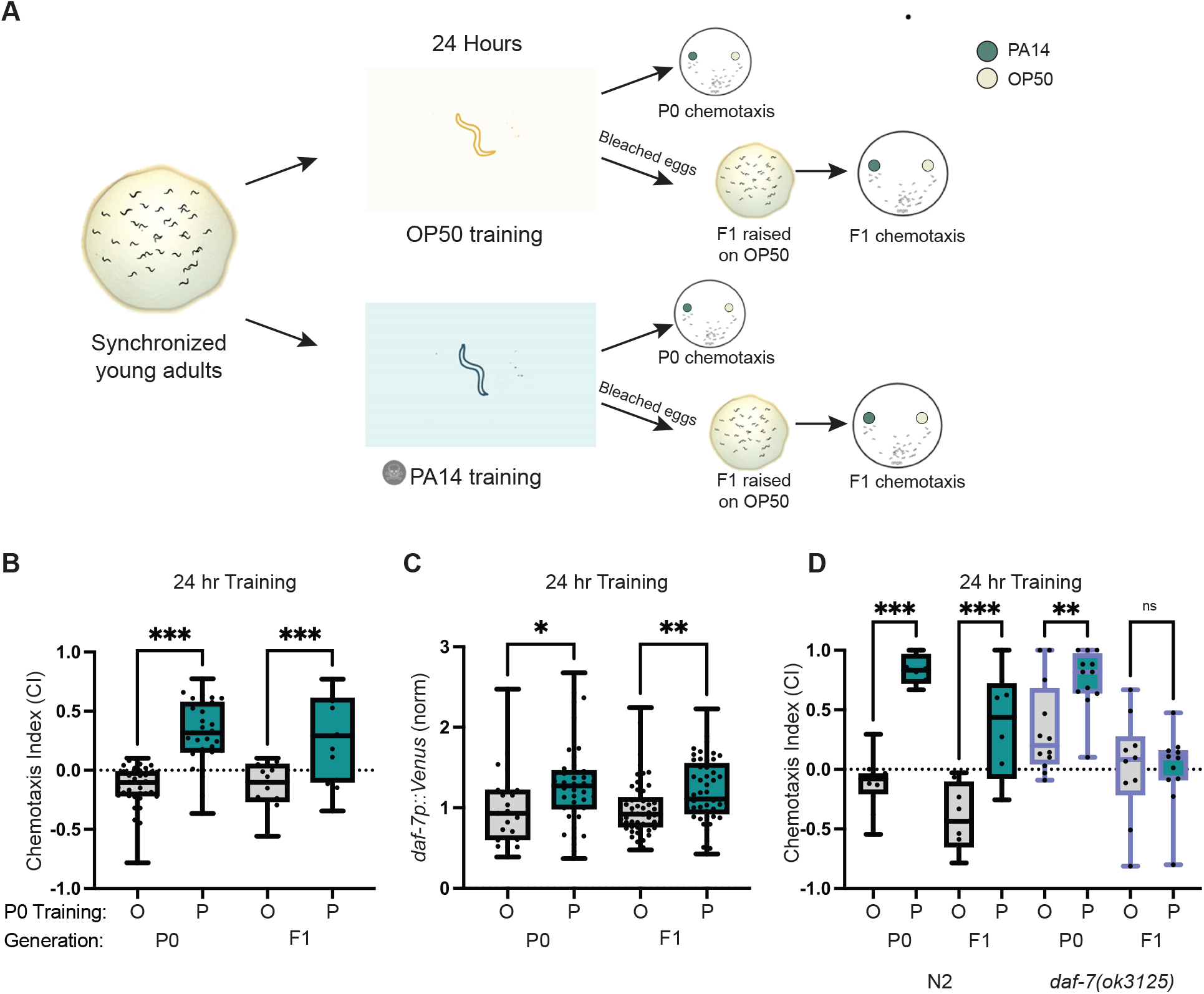
Intergenerational inheritance of pathogen avoidance in *C. elegans*. **A)** Experimental paradigm for training *C. elegans* on either *E. coli* OP50 or *P. aeruginosa* PA14 and assessing pathogen avoidance behavior across generations. Synchronized young adult worms were exposed to either OP50 (light background) or PA14 (blue background) for 24 hours. Following training, chemotaxis assays were conducted on the parental (P0) generation to assess their preference for OP50 or PA14. To examine transgenerational effects, P0 worms were bleached to obtain F1 embryos, which were then raised on standard OP50 plates until adulthood. Once matured, F1 worms from both OP50- and PA14-trained parents underwent chemotaxis assays to assess inherited pathogen avoidance. **B)** Chemotaxis results for young adults (P0) after 24 hour training on OP50 (O) or PA14 (P). F1 chemotaxis results are for animals descended from parents trained on OP50 or PA14. **C)** *daf-7p::Venus* expression in ASI neurons in both P0 and F1 after the training described in **A**). All values normalized to median of N2 OP50 trained animals. **D)** Chemotaxis results for N2 and *daf-7(ok3125)* mutants from behavioral paradigm in **A**). CI = ((# OP50) – (# PA14)) / Total. Wilxocon rank test, *p<0.05, **p<0.01, ***p<0.001; ns, not significant.

Although *daf-7* expression correlates with PA14 expression, prior research has shown that it is dispensable for PA14 avoidance in the P0 generation, but necessary for F1 avoidance (3–5). We, confirmed this to be true in our experimental paradigm as well; *daf-7* P0 animals learned to avoid PA14, but the no avoidance was observed in the F1 generation (**Fig. 1D**). Together, this establishes three reproducible observations: 1) F1 animals from PA14-trained P0 adults avoid PA14, 2) *daf-7* expression increases in PA14-trained adults, and their F1 progeny, and 3) *daf-7* is needed for inherited avoidance of PA14 in F1 animals.

### Germline loss uncovers skn-1b’s suppressive role of *daf-7* in pathogen avoidance behavior

Since parental exposure influences F1 behavior, we initially sought to investigate whether the germline influenced parental behavior. Germline loss is known to promote longevity in worms by reallocating energetic resources from reproduction to self-maintenance (15). During aging, germline-deficient animals exhibit increased resilience to heat and proteotoxic stress(16), but their response to acute stressors, such as pathogen exposure, remains poorly understood. To investigate this, we analyzed *glp-4*(*bn2*) temperature-sensitive (ts) mutants, which lack germ stem cell (GSC) proliferation due to a defect in valyl-tRNA synthetase(17). Intriguingly, germline-deficient mutants exhibited a two-fold resistance to PA14 compared to WT worms (**Fig. 2A**). This is consistent with the longevity-inducing effects of germline absence. Germline-deficient mutants accumulate excess lipids in the body wall and intestine, as yolk lacks a deposition site in the absence of eggs. This lipid overload activates SKN-1, the mammalian ortholog of Nuclear factor erythroid 2-related factor 2 (Nrf2), which induces the expression of lipid-metabolizing genes that promote stress resistance and longevity (15). Notably, among the three *skn-1* isoforms, *skn-1b* is expressed exclusively in the ASI neurons where *daf-7* is also expressed (11).

**Figure 2:**
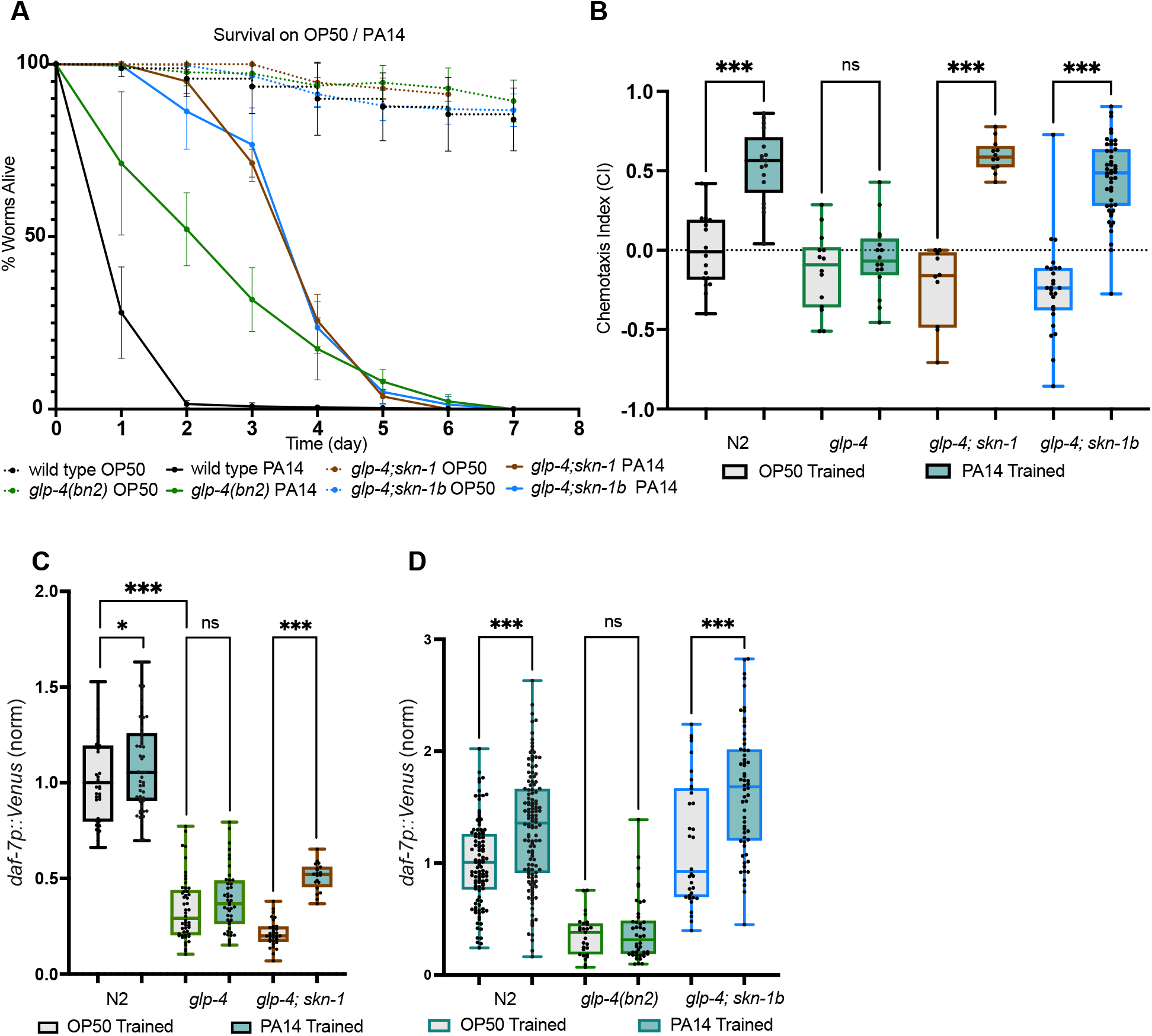
Germline Loss Uncovers SKN-1B’s Suppressive Role of DAF-7 and Pathogen Avoidance Behavior. **A**) N2, *skn-1(zj15), glp-4(bn2ts)*, and *glp-4(bn2ts)*;*skn-1(zj15), glp-4(bn2ts)*;*skn-1b(tm4241)* double mutants were assayed for lifespan at 25°C during exposure to OP50 (dashed lines) or PA14 (solid lines). **B**) Chemotaxis results for N2, *skn-1(zj15), glp-4(bn2ts), glp-4(bn2ts)*;*skn-1(zj15)*, and *glp-4(bn2ts)*;*skn-1b(tm4241)* trained on either OP50 (tan) or PA14 (green). **C**) *daf-7p::Venus* expression in the ASI neuron in N2, *skn-1(zj15), glp-4(bn2ts)*, and *glp-4(bn2ts)*;*skn-1(zj15)* animals. All values normalized to median of N2 OP50 trained animals. **D**) *daf-7p::Venus* expression in ASI in N2, *skn-1(zj15), glp-4(bn2ts)*, and *glp-4(bn2ts)*; *skn-1b(tm4241)* animals. All values normalized to median of N2 OP50 trained animals. CI = ((# OP50) – (# PA14)) / Total. Wilxocon rank test, *p<0.05, **p<0.01, ***p<0.001; ns, not significant.

Given *skn-1*’s role in longevity, we examined whether it contributes to survival in germline-deficient worms by testing *glp-4*(*bn2*); *skn-1*(*zj15*) and *glp-4*(*bn2*); *skn-1b*(*tm4241*) double mutants. The *skn-1*(*zj15*) allele is a hypomorph which disrupts *skn-1a* and *skn-1c* expression specifically, without disrupting *skn-1b* expression (18). Conversely, the *skn-1b*(*tm4241*) null allele deletes a large portion of an exon that is specific to the *skn-1b* isoform, and does not affect other isoforms (11). While *skn-1* is critical for longevity in germline-deficient mutants over long timescales on non-pathogenic bacteria (15), our results indicate that its loss does not significantly impact resilience under acute pathogenic stress (e.g., during the 8-day PA14 exposure) (**Fig. 2A**). This suggests that SKN-1’s role in the germline-deficient context is primarily linked to long-term survival rather than immediate pathogen resistance, prompting us to investigate whether it instead influences behavioral adaptation to PA14.

To test the role of *skn-1* on pathogen avoidance in germline-deficient mutants, we examined learned PA14 avoidance in *glp-4; skn-1* and *glp-4; skn-1b* double mutants. Germline-deficient worms failed to develop PA14 aversion, but in the *skn-1* or *skn-1b* mutant background, PA14 avoidance was restored (**Fig. 2B**). These results suggest that the loss of SKN-1, and specifically the SKN-1B isoform, is sufficient to restore pathogen avoidance behavior in germline deficient worms. We next investigated *daf-7* expression in germline deficient worms and found that *glp-4* mutants exhibited significantly lower baseline levels of *daf-7p::Venus* compared to wild-type.

However, loss of *skn-1* in the *glp-4* background restored a significant induction of *daf-7p::Venus* expression following PA14 exposure, despite the overall baseline remaining lower (**Fig. 2C**). This enhanced change in expression suggests that the loss of *skn-1* facilitates a more robust neuroendocrine response to infection in germline-deficient worms, potentially underpinning the restoration of pathogen avoidance behavior. Notably, the loss of the *skn-1b* isoform alone in the *glp-4* background was sufficient to enable PA14-induced expression of *daf-7p::Venus*, without significantly affecting baseline expression (**Fig. 2D**). This suggests that *skn-1* plays a critical role in modulating *daf-7* expression in response to pathogen exposure and its associated behavioral outcomes, in germline-deficient worms.

We next assessed the role of *skn-1* in a germline-intact background to determine whether its function extends beyond the germline-deficient context. Unlike the germless background, there was no significant difference in *daf-7p::Venus* expression between *skn-1* mutants and wild-type worms following PA14 exposure, for both the P0 and F1 generations (**Fig. 3A**). However, loss of the *skn-1b* isoform, led to elevated *daf-7p::Venus* expression in all conditions compared to wild-type (**Fig. 3B**). This elevated expression also appeared to abolish the differential expression of *daf-7p::Venus* in the PA14 P0 and F1 progeny (**Fig. 3B**).

**Figure 3:**
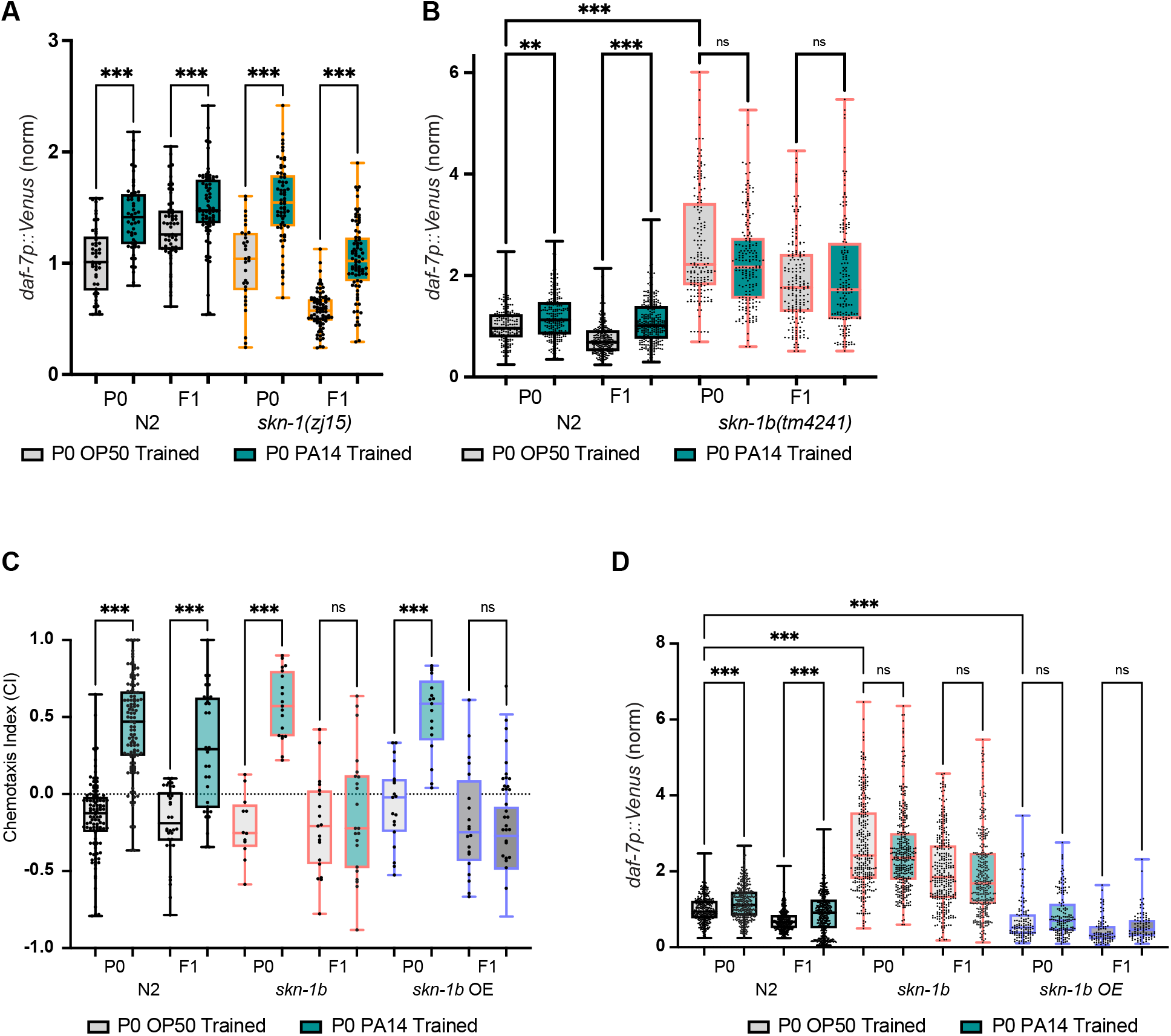
Loss or overexpression of *skn-1b* dysregulates the pathogen-induced daf-7 response and prevents inherited learning in F1 animals. **A**) *daf-7p::Venus* expression in the ASI neuron in N2 and *skn-1(zj15)*, animals in the P0 (trained) and F1 generations. All values normalized to median of N2 OP50 trained animals. **B**) *daf-7p::Venus* expression in the ASI neuron in N2 and *skn-1b(tm4241)* animals in the P0 (trained) and F1 generations. All values normalized to median of N2 OP50 trained animals. **C**) Chemotaxis of OP50 and PA14-trained P0 animals, and their F1 progeny, in N2, *skn-1b(tm4241)* and *skn-1b* overexpression (OE) backgrounds. **D**) *daf-7p::Venus* expression in the ASI neuron in N2, *skn-1b(tm4241)*, and *skn-1b* overexpression (OE) backgrounds for P0 (trained) and F1 generations. All values normalized to median of N2 OP50 trained animals. CI = ((# OP50) – (# PA14)) / Total. Wilxocon rank test, *p<0.05, **p<0.01, ***p<0.001; ns, not significant.

Given that the loss of *skn-1b* was sufficient to increase *daf-7p::Venus* expression, we next investigated whether overexpression of *skn-1b* would suppress *daf-7p::Venus* expression, and whether this would have an impact on PA14 avoidance in P0 and F1. To this end, we generated an extrachromosomal array *skn-1bp::aid::skn-1b::SL2::aid::mCherry* to overexpress *skn-1b* under the *skn-1b* promoter. As expected, *daf-7p::Venus* expression was significantly lower in the *skn-1b* overexpression background compared to control (**Fig. 3D**). In addition, the overexpression of *skn-1b* appeared to suppress induction of *daf-7p::Venus* as a result of PA14 exposure. However, this suppression did not appear to affect the chemotaxis of the P0 generation, but it did suppress the learning of the F1 generation (**Fig. 3C**). This was similar to what was observed with the loss of *skn-1b*: the dysregulation of the *daf-7* response in the P0 generation does not affect learned PA14 avoidance, but it does affect F1-inherited avoidance (**Fig. 1D**). Over-expression of *skn-1b* suppressed *daf-7* expression, and loss of *skn-1b* increased *daf-7* expression. Both of these perturbations eliminated PA14-induced changes in *daf-7* expression, which did not seem to affect P0 chemotaxis. However, loss of PA14-induced induction of *daf-7* did eliminate F1 PA14 avoidance. These findings indicate that SKN-1B negatively regulates *daf-7* expression, and dysregulation of *daf-7* expression via loss or over-expression of *skn-1b* disrupts the *daf-7* response to PA14, which is necessary for F1-acquired pathogen avoidance.

### Parental SKN-1B is necessary for inherited PA-14 avoidance in F1 animals

SKN-1B’s regulatory effect on *daf-7* expression appears to be important for the F1 generation, but in both of the prior perturbations, SKN-1B activity was perturbed in both P0 and F1 animals. To determine whether SKN-1B is needed in the P0 or F1 generation for F1 pathogen avoidance behavior, we utilized the auxin-inducible degradation (AID) system. This system enables precise, conditional protein depletion using a set of single-copy, tissue-specific TIR1-expressing strains integrated at well-characterized genetic loci, allowing targeted degradation of AID-tagged proteins in specific tissues without interfering with fluorescent reporters (19). To assess the generational role of *skn-1b* on *daf-7* expression and pathogen avoidance, we tested two different conditions. In the P0-null condition, P0 animals were trained on plates of OP50 or PA14 that were supplemented with or without auxin. Since auxin-induced degradation requires less than an hour of exposure (19), and training is for 24 hours, P0 animals were trained in the absence of *skn-1b* for the majority of their training period. However, bleached eggs from this generation were raised on auxin-free plates of OP50 for their entire development to adulthood (3 days) so that adult F1 *skn-1*b expression was not degraded due to auxin exposure. In the F1-null condition, F1 (but not P0) animals were trained in the presence of auxin. In this condition, only the F1 animals experience a depletion of SKN-1B.

In the P0-null context, auxin-induced *skn-1b* depletion during OP50 training led to an increase in *daf-7p::Venus* expression in the P0 generation **(Fig. 4A)**. However, *daf-7* expression was not increased due to PA14 exposure in the *daf-7*-elevated context. This was consistent with observations made in the *skn-1b*(*tm4241*) background: loss of *skn-1b* leads to elevated levels of *daf-7* expression that are not increased due to PA14 exposure. Even though auxin-induced degradation occurred during parental training, the elevated *daf-7* expression was also observed in the adult F1 generation (**Fig. 4A**). Like the P0 generation, the elevated baseline expression of *daf-7* prevented a further increase in the F1 generation descended from PA14-trained adults. This indicates that the loss of SKN-1B in the P0 generation elevates *daf-7* expression in both the P0 and F1 generations. In turn, PA14-induced increases in *daf-7* expression are not observed when baseline *daf-7* expression is elevated.

**Figure 4:**
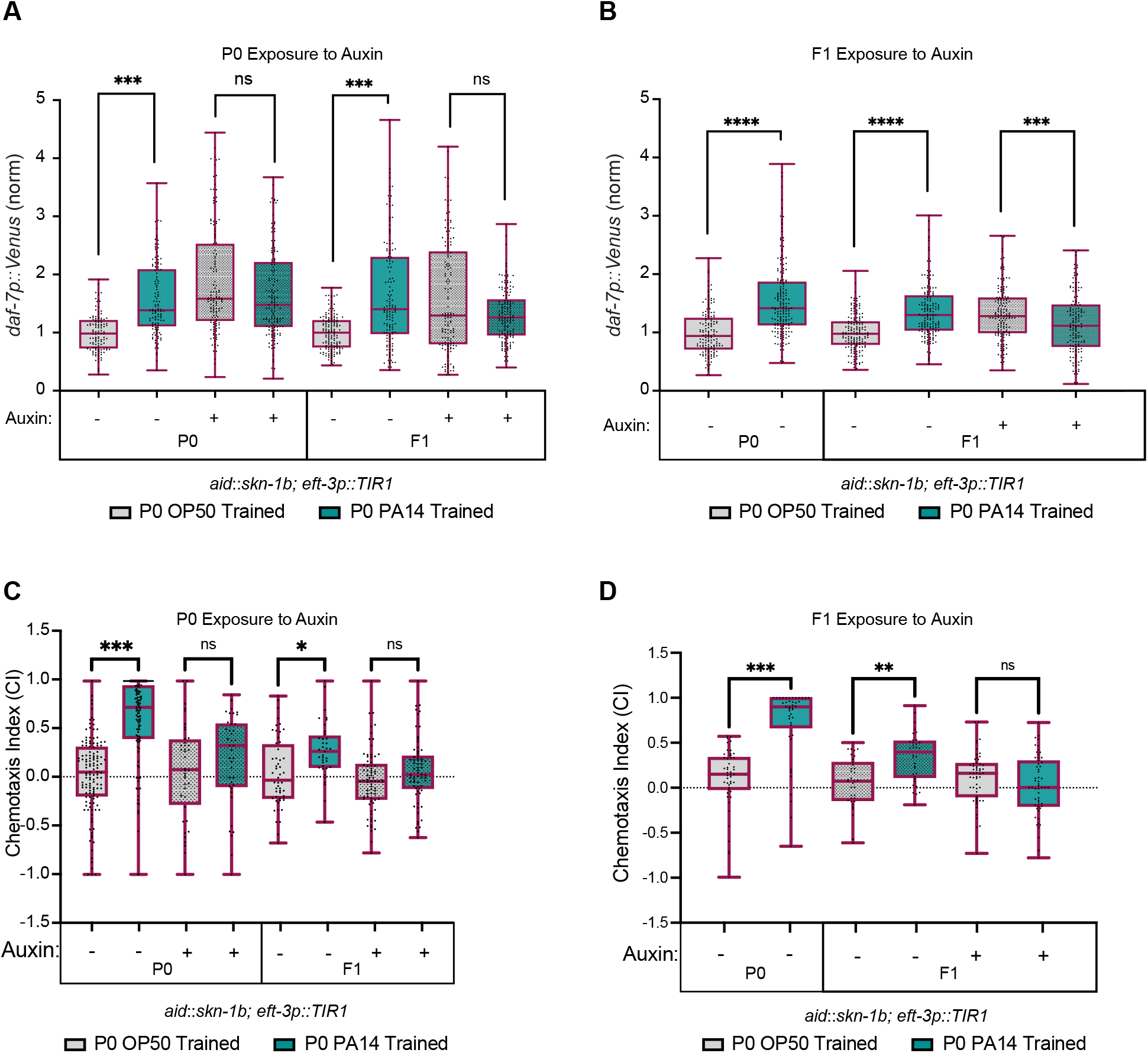
SKN-1B is needed in both the P0 and F1 generations for inherited PA14 avoidance in F1. **A**) *daf-7p::Venus* expression in the ASI neuron in aid-tagged *skn-1b* P0 (trained) and F1 animals. Auxin exposure (− or +) occurred during the 24 hour training in the P0 generation. All values normalized to median of N2 OP50 trained animals. **B**) *daf-7p::Venus* expression in the ASI neuron in aid-tagged *skn-1b* P0 (trained) and F1 animals. Auxin exposure (− or +) occurred during the 24 hour period before chemotaxis in the F1 generation. All values normalized to median of N2 OP50 trained animals. **C**) Chemotaxis of OP50 and PA14-trained P0 animals, and their F1 progeny, in the aid-tagged *skn-1b* background. Auxin exposure (− or +) occurred during the 24 hour training in the P0 generation. **D**) Chemotaxis of OP50 and PA14-trained P0 animals, and their F1 progeny, in the aid-tagged *skn-1b* background. Auxin exposure (− or +) occurred during the 24 hour period before chemotaxis in the F1 generation. CI = ((# OP50) – (# PA14)) / Total. Wilxocon rank test, *p<0.05, **p<0.01, ***p<0.001; ns, not significant.

If SKN-1B was depleted in the F1 generation, a similar increase in *daf-7p::Venus* expression was observed which suppressed a further increase in *daf-7p::Venus* expression due to parental PA14 training (**Fig. 4B**). In fact, the F1 generation from PA14-trained P0 appeared to have slightly less *daf-7p::Venus* expression when compared to the F1s descended from OP50-trained P0s. A slight decrease in median expression in the P0-null background was also observed in the PA14-trained context, but this difference was not significant.

In the *skn-1b* mutant, *skn-1b* overexpression, and *daf-7* mutant backgrounds, chronic disruption of *daf-7* expression did not affect P0 learned avoidance of PA14, but it did affect the acquired avoidance in the F1 generation (**Fig. 1 – 3**). However, *acute* depletion of SKN-1B during P0 training *did* appear to prevent learned PA14 avoidance in the P0 generation (**Fig. 4C**). This loss of PA14 avoidance was also observed in the F1 generation. Acute depletion of SKN-1B in the F1 generation also led to a loss of F1-inherited PA14 avoidance (**Fig. 4D)**. These results reveal that SKN-1B is needed in the P0 generation to influence F1 PA14 avoidance, but that it is also needed in the F1 generation as well. Curiously, while chronic disruption of *skn-1b* and/or *daf-7* does not affect learned PA14 avoidance in the P0 generation, an acute disruption of *skn-1b* and *daf-7* does affect learned pathogen avoidance.

## Discussion

The findings presented in this study provide evidence for the role of SKN-1B and its influence on DAF-7 signaling in mediating intergenerational pathogen avoidance behavior in *C. elegans*. Our data confirm previous observations that 24 hours of exposure to the pathogenic bacterium *P. aeruginosa* PA14 robustly elevates *daf-7* expression in the ASI neurons of the parental (P0) generation, and that this elevated expression is maintained in the F1 progeny, even though the progeny themselves have not been directly exposed to PA14. We also confirm that *daf-7(ok3125)* mutants can acquire pathogen avoidance behavior after training but fail to transmit this behavior to the F1 generation, underscoring the necessity of DAF-7 signaling for intergenerational inheritance of this learned response (3, 13).

Further experimentation examining the role of the germline revealed that germline-deficient mutants (*glp-4*(*bn2*)) exhibit extended resistance to PA14, consistent with prior reports linking germline loss to increased longevity. These worms not only survived 2-to 3-fold longer when challenged with PA14, but also failed to exhibit the typical avoidance response after 24 hours of exposure to PA14. Previous studies conducted under OP50 feeding conditions over the course of 30 days have shown that loss of *skn-1* diminishes the extended longevity observed in germline-deficient mutants(15). In addition to this, pathogen exposure is known to induce *skn-1* expression in several tissues, which in turn confer increased resistance to pathogen infection (6–9).

However, in our PA14 longevity assays, which monitor survival over an 8-day acute pathogenic exposure, the loss of *skn-1* or the *skn-1b* isoform did not abrogate the longevity extension in the *glp-4* background. Intriguingly, the loss of *skn-1*, and specifically the *skn-1b* isoform, in the *glp-4* background did restore PA14 avoidance after PA14-training. This indicates that the loss of PA14 avoidance in the *glp-4* background was not necessarily due to increased longevity, since the *skn-1;glp-4* worms still maintained extended longevity on PA14.

In the *glp-*4 background, baseline *daf-7* expression was significantly lower than that observed in N2, and no appreciable difference was detected between OP50- and PA14-trained conditions. However, in *glp-4* mutants carrying hypomorphic *skn-1*(*zj15*), differential *daf-7* expression levels between OP50- and PA14-trained conditions were restored, even though overall *daf-7* expression remained lower than N2. Notably, the loss of the *skn-1b* isoform alone in the *glp-4* background was sufficient to not only restore learning and *daf-7* differential expression, but it also restored *daf-7* baseline levels. These results suggest that the induction of pathogen avoidance behavior in a germless background may depend on a sufficient differential in *daf-7* expression to serve as the trigger for avoidance responses. However, it is important to note that in *daf-7* animals, which do not lack a germline, *daf-7* did *not* appear to be necessary for P0 PA14 learned avoidance, but *daf-7* was necessary for the F1 acquired avoidance.

For N2 animals with intact germlines, the *skn-1(zj15)* hypomorph did not significantly alter *daf-7* expression compared to wild type, whereas selective deletion of the *skn-1b* isoform resulted in markedly elevated *daf-7* levels under all conditions. The elevated daf-7 appeared to prevent further increases in daf-7 expression due to PA14 exposure. Unlike the *glp-4* germless mutant, the loss of a PA14-induced increase of *daf-7* expression did not affect the learned avoidance of PA14 in the P0 generation, but the avoidance was lost in the F1 generation. The converse also appeared to be true: an overexpression of *skn-1b* dropped *daf-7* levels and also inhibited PA14-induced increases in *daf-7* expression. As in the *skn-1b* mutant, loss of PA14-induced regulation of *daf-7* (due to elevated *skn-1b* expression) did not affect P0 learned avoidance, but it did affect F1 acquired avoidance. Given the colocalization of SKN-1B and DAF-7 in the ASI neurons, the inverse relationship shown implies that SKN-1B modulates *daf-7* expression, ultimately influencing pathogen avoidance behavior.

ChIP-seq and RNAseq analysis of a *skn-1* gain-of-function allele, *skn-1gf*(*lax188*), has revealed several genes regulated by *skn-1*, but *daf-7* has not appeared in these data (7, 8, 10, 20, 21). However, the *skn-1gf*(*lax188*) allele does not affect *skn-1b* expression, so most of the genes detected in these studies are likely from *skn-1a/c* activity (10). Interestingly, even though these alleles are expressed in the intestine in addition to ASI, the activation of these alleles in ASI is sufficient to change the proteostatic balance within intestinal cells. PA14 has been shown to induce the expression and nuclear localization of *skn-1* in intestinal cells, with no apparent change in the ASI neurons. However, these observations are primarily the result of the *skn-1c* isoform (8). Our data suggest that despite activation of *skn-1c* in intestinal cells, PA14 may inhibit the activity of the *skn-1b* isoform, leading to an increase of *daf-7* expression. This activity affects learned PA14 avoidance in germless mutants, but not in animals with intact germlines. However, the F1 progeny of these animals do rely on PA14-induced changes to *skn-1b* and *daf-7* activity.

We found that SKN-1B is necessary in both the P0 and F1 generations to enable changes in *daf-7* expression and acquired PA14 avoidance in F1 animals. The loss of SKN-1B in either generation eliminates the PA14-induced elevation of *daf-7* expression and PA14 avoidance in the F1 generation. Curiously, acute loss of SKN-1B during PA14 training also reduced learned avoidance in the P0 generation. This was surprising since the *skn-1b* mutant had no effect on P0 learning. The chronic absence of *skn-1b* may lead to compensatory mechanisms to enable PA14 learned avoidance, while the acute loss of SKN-1B during the training period may be too short to allow for this to occur.

How *skn-1b* is directly influenced by PA14, and how its changes to *daf-7* expression influence the F1 generation, remain to be answered. The activation of *skn-1c* in ASI has systemic effects on gene expression throughout the animal (10). DAF-7 has been shown to play a critical role in transgenerational avoidance of PA14 (3), but in the context we explored here, transgenerational inheritance of PA14 avoidance was not observed. Since our observations only span one generation, the molecular mechanisms for the single-generation inheritance of pathogen avoidance need not rely on epigenetic mechanisms (3–5). Instead, PA14-induced changes to *skn-1b* and *daf-7* likely alter *in utero* signaling during embryo development. This signaling is likely absent in the F1 generation which has not been exposed to PA14, and hence we do not observe learned PA14-avoidance in the F2 generation. The observations made here may work in parallel to transgenerational effects that rely on epigenetic mechanisms (3–5, 13).

## Materials and Methods

### Worm Maintenance and Preparation for Imaging

Worms were grown and maintained under standard conditions using 6 cm nematode growth medium (NGM) plates and fed *Escherichia coli* OP50 bacteria (22).

### All worms were grown at □20°C on 6cm Nematode Growth Medium (NGM) agar plates and seeded with with 250 □μL *E. coli* strain OP50

#### CRISPR/Cas9 knock-in strategy

The AID-GSGGG-3xFLAG-skn-1bN2 strain was generated using CRISPR/Cas9-mediated genome editing, performed by SunnyBiotech Co. Ltd. This strain was designed to introduce an auxin-inducible degron (AID) tag, a GSGGG flexible linker, and a 3×FLAG epitope tag at the N-terminus of the endogenous *skn-1b* locus in the N2 wild-type background. To achieve this, a homology-directed repair (HDR) approach was used, in which a guide RNA (sgRNA) targeting the *skn-1b* N-terminal coding sequence was co-injected with the Cas9 protein to induce a site-specific double-strand break. A synthetic single-stranded DNA oligonucleotide (ssODN) repair template containing the AID-GSGGG-3xFLAG sequence flanked by ∼35-50 base pair homology arms was provided to facilitate HDR-mediated knock-in.

Following microinjection, F1 progeny were screened for successful insertion using PCR genotyping with primers flanking the targeted integration site. Positive strains were confirmed via Sanger sequencing to verify seamless in-frame integration of the AID-GSGGG-3xFLAG sequence without additional mutations. The successfully edited strain was maintained under standard *C. elegans* culture conditions at 20°C on nematode growth medium (NGM) plates seeded with E. coli OP50.

### Generation of transgenic strains

The *skn-1bp*::AID::SKN-1B::SL2::AID::mCherry transgenic strain was generated using Gibson assembly to construct the transgene, followed by microinjection into *C. elegans* for extrachromosomal array formation. This construct was designed to express AID-tagged *skn-1b* under the control of the endogenous *skn-1b* promoter, followed by an SL2 spliced leader sequence allowing co-expression of AID-tagged mCherry as a fluorescent reporter within the same transcript. This bicistronic design ensured that SKN-1B expression could be visualized and tracked *in vivo* via mCherry fluorescence. Successfully transformed animals were identified based on neuronal RFP fluorescence and further validated by PCR genotyping to confirm the presence of the transgene. The expression of the mCherry proteins was confirmed using fluorescence microscopy, ensuring proper localization and co-expression within the targeted neuronal cells. The transgenic strain was subsequently maintained by selecting for RFP-positive animals at each generation.

#### Synchronization and pathogen training

To assess avoidance behavior toward *Pseudomonas aeruginosa* strain PA14, *C. elegans* were synchronized and subjected to pathogen training. Synchronization was initiated by placing 20– 30 young adults on lawns of the non-pathogenic *Escherichia coli* strain OP50, allowing them to lay eggs for 24 hours. After this period, adult worms were removed, leaving the eggs to hatch and develop. The eggs were incubated at 20°C for three days, reaching adulthood by day three. For pathogen training, young adult worms were transferred onto either full lawns of PA14 (for the experimental group) or OP50 (for the control group) for 24 hours. Full lawns were prepared three days prior to training. A single colony of either PA14 or OP50 was inoculated into 3 mL of LB medium and grown overnight on a floor shaker at 37°C. The cultures typically reached an optical density (OD) of approximately 2.5. Before plating, the cultures were diluted to an OD of 1.0 with Lorea Broth (LB). Subsequently, 400 µL of the adjusted culture was spread evenly onto 6 cm nematode growth media (NGM) plates. The plates were incubated at room temperature for one day, followed by an additional day at 27°C to ensure the virulence of PA14 and readiness of OP50 lawns for training.

#### Chemotaxis Assay

To quantify avoidance behavior following OP50 or PA14 training, a chemotaxis assay was conducted. Chemotaxis plates were prepared by inoculating samples of OP50 and PA14 overnight, and then diluting each sample to an optical density (OD) of 1. Then 20 µL of OP50 and 20ul of PA14 were spotted onto an NGM plate equidistant from each other. Approximately 100 trained worms were collected from training plates using M9 buffer and transferred into microcentrifuge tubes. Worms were pelleted by gravity, and the supernatant was removed and replaced with fresh M9 buffer. This wash step was repeated three times before worms were dispensed for chemotaxis assay. Approximately 50 worms (10ul) were aliquoted using a glass Pasteur pipette onto the origin spot of the chemotaxis plate, and excess liquid was removed with a 10ul pipette to enable rapid foraging. Worms were allowed to move freely for one hour, after which sodium azide was added to the bacterial spots to paralyze the worms that reached either spot.

The chemotaxis index (CI) was calculated as:

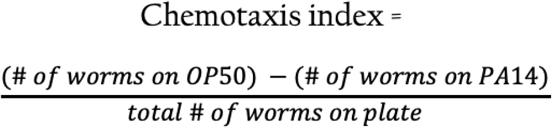

A positive CI indicates a preference for OP50, while a negative CI suggests attraction to PA14. This index served as a measure of avoidance behavior, with a higher CI indicating stronger avoidance of PA14.

#### Microscopy and imaging of neuronal expression

To verify neuronal expression of Venus and mCherry reporters linked to *daf-7* and *skn-1b* expression respectively, laser scanning microscopy was employed. Worms washed from training plates with M9, and pelleted by gravity, and the supernatant was removed and replaced with fresh M9 buffer. This wash step was repeated three times before worms were placed on imaging pads. Once placed on 2% agarose pads, worms were arranged head-to-tail with an eyelash pick and imaged using the LSM-700 laser scanning confocal microscope (Zeiss). Z-stacks were taken of the ASI neurons, and the images were tiled and stitched to generate a maximum projection. Regions of interest (ROIs) were manually circled in ImageJ, and the average fluorescence intensity for each condition was measured. Background subtraction was not applied, as background noise was consistently low across all samples due to the thorough three-round washing procedure. Fluorescence values from this microscopy analysis were used to quantify the expression of *daf-7* and *skn-1b* in the ASI neuron under different training conditions.

#### Survival Assays

To obtain synchronized populations while minimizing potential developmental defects from bleaching, adult *C. elegans* were allowed to lay eggs on OP50-seeded plates for 24 hours. The following day, adults were washed off, leaving only eggs. These eggs developed into adults over approximately three days and were then transferred to full-lawn pathogen training plates containing either *E. coli* OP50 or *P. aeruginosa* PA14, with 100 worms per plate. After 24 hours, surviving worms were transferred to fresh OP50 or PA14 plates. This process was repeated daily until all worms had died, with survival recorded each day. Survival was assessed by whether worms moved when prodded. The assay was performed in three biological replicates.

#### Auxin Inducible Degradation

To selectively degrade SKN-1B, the auxin-inducible degradation (AID) system was employed using NGM plates supplemented with 400 µM potassium auxin (KAA), a water-soluble auxin analog that is non-photodegradable. Auxin plates were prepared by adding 400 µM KAA (PhytoTech Labs) from a 100 mM stock solution dissolved in water to autoclaved NGM agar after cooling to ∼55°C. Plates were left to dry in the dark overnight before being seeded with E. coli OP50 or P. aeruginosa PA14.

To induce SKN-1B degradation, worms carrying the AID-tagged SKN-1B transgene and a ubiquitously expressed TIR1 transgene (Peft-3::TIR1::BFP) were transferred to NGM + 400 µM KAA plates under two distinct experimental conditions: (1) P0 worms were placed on auxin-containing plates during training on either OP50 or PA14, or (2) only their progeny (F1 generation) were exposed to auxin, while the P0 generation remained on standard NGM plates. The *Peft-3::TIR1::BFP* transgene, which expresses BFP as an internal control, was used to assess auxin-dependent protein degradation. Since the construct couples TIR1 expression to BFP, auxin-induced degradation of SKN-1B should correlate with a reduction in BFP fluorescence.

#### Statistical Analysis

Data from chemotaxis and fluorescence assays were analyzed using the Kruskal-Wallis test to account for multiple genotypes and both *E. coli* OP50 and *P. aeruginosa* PA14 conditions.

Wilcoxon tests were performed *post hoc*, with Sidak correction for multiple comparisons. Given the non-normal distribution of data and differences in standard deviations across groups, this non-parametric approach allowed for robust comparisons across multiple experimental conditions. Post-hoc tests were conducted where necessary to identify significant pairwise differences. Each experiment was performed with at least three biological replicates to ensure reproducibility and statistical reliability.

#### Experimental models: organisms/strains

Worms were cultured according as previously described, and maintained at 20°C unless otherwise indicated. The following strains were used: N2 CGC hermaphrodite stock, GA1058: *skn-1b(tm4241)*, QL196: *drcSI61[unc-119;Ptph-1::mCherry] I; drcSI7[unc-119; daf-7p::Venus] II*, SS104: *glp-4(bn2)*, QV225: *skn-1(zj15)*, AGG0153: *glp-4(bn2);skn-1(zj15); drcSI7[unc-119; daf-7p::Venus]*, AGG0164: *glp-4(bn2);skn-1b(tm4241); drcSI7[unc-119; daf-7p::Venus]*, RB2302: *daf-7(ok3125)*, AGG0165: PHX796 *AID::skn-1b::3x-FLAG (syb7960);*JDW225 *eft-3p::TIR1::F2A::mTagBFP2::AID*::NLS::tbb-2 3’UTR; QL196 drcSI7[unc-119; daf-7p::Venus]*, AGG0166: SS104 *glp-4(bn2);* PHX796 *AID::skn-1b::3x-FLAG (syb7960);*JDW225 *eft-3p::TIR1::F2A::mTagBFP2::AID* ::NLS::tbb-2 3’UTR; QL196 drcSI7[unc-119; daf-7p::Venus]*

## Author Contributions

R.P. and A.G. conceived the experiments. R.P., M.B., and E.O. performed chemotaxis and training experiments. R.P. performed imaging experiments. R.P. and A.G. analyzed and wrote the manuscript.

## Acknowledgements

We would like to thank J. Alonzo, I. Mosley, P. Jackson, and N. Kose for their assistance in preparing reagents for many of the assays in the experiments. We thank J. Kim, G. Seydoux, and Y. Kim for helpful discussions and suggestions. We also thank Q. Ch’ng for strain QL196, and J. Tullet for strain GA1058. Some strains were provided by the CGC, which is funded by NIH Office of Research Infrastructure Programs (P40 OD010440). A.G. acknowledges funding from NIH (R35GM124883).

---

**Supplemental Figure 1: Inherited PA14 avoidance observed in the F1, but not subsequent, generations. A**) 24 h of parental training on PA14 induces pathogen avoidance in the parents (P0) and progeny (F1), but not in the subsequent generations. CI = ((# OP50) – (# PA14)) / Total. Wilxocon rank test, *p<0.05, **p<0.01, ***p<0.001; ns, not significant.

## References

1. F. Zhang, et al., Caenorhabditis elegans as a Model for Microbiome Research. Frontiers in Microbiology 8, 485 (2017).

2. B. S. Samuel, H. Rowedder, C. Braendle, M.-A. Félix, G. Ruvkun, Caenorhabditis elegans responses to bacteria from its natural habitats. Proceedings of the National Academy of Sciences 113, E3941–E3949 (2016).

3. R. S. Moore, R. Kaletsky, C. T. Murphy, Piwi/PRG-1 Argonaute and TGF-β Mediate Transgenerational Learned Pathogenic Avoidance. Cell 177, 1827-1841.e12 (2019).

4. R. Kaletsky, et al., Molecular Requirements for <em>C. elegans</em> Transgenerational Epigenetic Inheritance of Pathogen Avoidance. bioRxiv 2025.01.21.634111 (2025). 10.1101/2025.01.21.634111.

5. D. P. Gainey, A. V. Shubin, C. P. Hunter, Irreproducibility of transgenerational learned pathogen-aversion response in C. elegans. (2024). 10.1101/2024.06.01.596941.

6. D. ana Papp, P. eter Csermely, C. Sőti, A Role for SKN-1/Nrf in Pathogen Resistance and Immunosenescence in Caenorhabditis elegans. PLOS Pathogens 8, e1002673 (2012).

7. J. D. Nhan, et al., Redirection of SKN-1 abates the negative metabolic outcomes of a perceived pathogen infection. Proceedings of the National Academy of Sciences 116, 22322–22330 (2019).

8. C. Gabald\’ on O. Karakuzu, D. A. Garsin, SKN-1 activation during infection of Caenorhabditis elegans requires CDC-48 and endoplasmic reticulum proteostasis. Genetics 228, iyae131 (2024).

9. R. van der Hoeven, K. C. McCallum, M. R. Cruz, D. A. Garsin, Ce-Duox1/BLI-3 Generated Reactive Oxygen Species Trigger Protective SKN-1 Activity via p38 MAPK Signaling during Infection in C. elegans. PLOS Pathogens 7, e1002453 (2011).

10. C. D. Turner, N. L. Stuhr, C. M. Ramos, B. T. Van Camp, S. P. Curran, A dicer-related helicase opposes the age-related pathology from SKN-1 activation in ASI neurons. Proceedings of the National Academy of Sciences 120, e2308565120 (2023).

11. N. Tataridas-Pallas, et al., Neuronal SKN-1B modulates nutritional signalling pathways and mitochondrial networks to control satiety. PLOS Genetics 17, e1009358 (2021).

12. J. D. Meisel, O. Panda, P. Mahanti, F. C. Schroeder, D. H. Kim, Chemosensation of bacterial secondary metabolites modulates neuroendocrine signaling and behavior of C. elegans. Cell 159, 267–280 (2014).

13. A. G. Pereira, X. Gracida, K. Kagias, Y. Zhang, C. elegans aversive olfactory learning generates diverse intergenerational effects. Journal of Neurogenetics 34, 378–388 (2020).

14. Y. Zhang, H. Lu, C. I. Bargmann, Pathogenic bacteria induce aversive olfactory learning in Caenorhabditis elegans. Nature 438, 179–184 (2005).

15. M. J. Steinbaugh, et al., Lipid-mediated regulation of SKN-1/Nrf in response to germ cell absence. eLife 4, e07836 (2015).

16. A. Ben-Zvi, E. A. Miller, R. I. Morimoto, Collapse of proteostasis represents an early molecular event in Caenorhabditis elegans aging. Proc Natl Acad Sci U S A 106, 14914– 14919 (2009).

17. S. Rastogi, et al., Caenorhabditis elegans glp-4 Encodes a Valyl Aminoacyl tRNA Synthetase. G3 Genes|Genomes|Genetics 5, 2719–2728 (2015).

18. L. Tang, W. Dodd, K. Choe, Isolation of a Hypomorphic skn-1 Allele That Does Not Require a Balancer for Maintenance. G3 (Bethesda) 6, 551–558 (2015).

19. G. E. Ashley, et al., An expanded auxin-inducible degron toolkit for Caenorhabditis elegans. Genetics 217, iyab006 (2021).

20. T. Nair, B. A. Weathers, N. L. Stuhr, J. D. Nhan, S. P. Curran, Serotonin deficiency from constitutive SKN-1 activation drives pathogen apathy. Nature Communications 15, 8129 (2024).

21. C. M. Ramos, S. P. Curran, Comparative analysis of the molecular and physiological consequences of constitutive SKN-1 activation. GeroScience 45, 3359–3370 (2023).

22. S. Brenner, The genetics of Caenorhabditis elegans. Genetics 77, 71–94 (1974).

