## Supplementary figures and images for "The Maternal Effect of SKN-1B and DAF-7 on Intergenerational Pathogen Avoidance Learning in *C. elegans*"

### Supplementary Figure 1

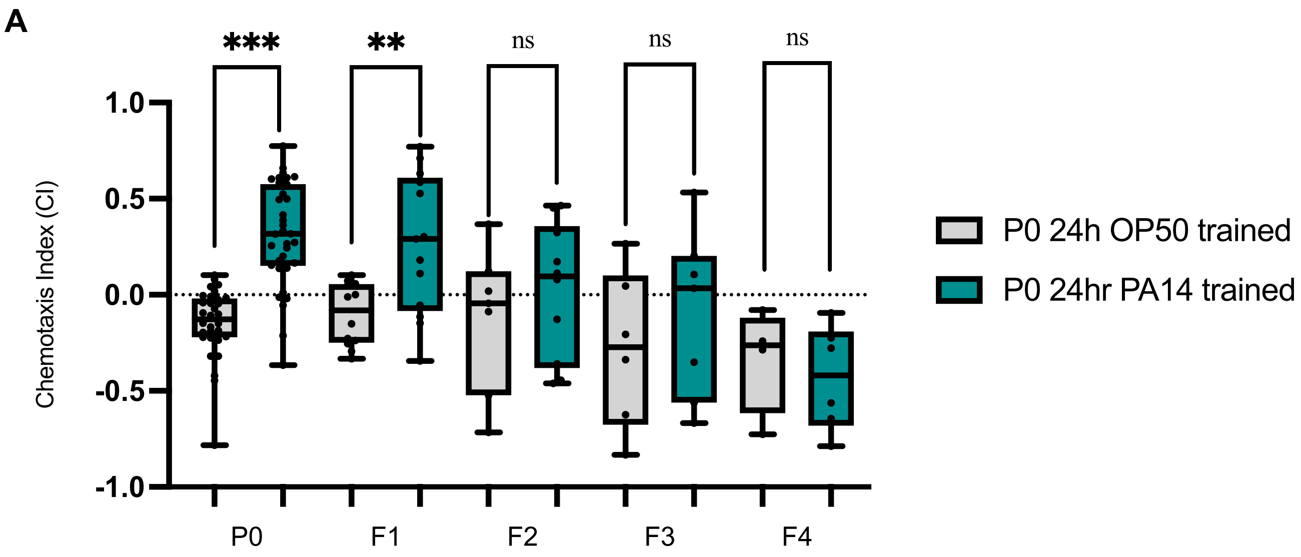
